# Comprehensive synthetic genetic array analysis of alleles that interact with mutation of the *Saccharomyces cerevisiae* RecQ helicases Hrq1 and Sgs1

**DOI:** 10.1101/2020.08.28.272666

**Authors:** Elsbeth Sanders, Phoebe A. Nguyen, Cody M. Rogers, Matthew L. Bochman

**Affiliations:** Molecular and Cellular Biochemistry Department, Indiana University, Bloomington, IN 47405

**Keywords:** *Saccharomyces cerevisiae*, *HRQ1*, *SGS1*, DNA helicase, yeast

## Abstract

Most eukaryotic genomes encode multiple RecQ family helicases, including five such enzymes in humans. For many years, the yeast *Saccharomyces cerevisiae* was considered unusual in that it only contained a single RecQ helicase, named Sgs1. However, it has recently been discovered that a second RecQ helicase, called Hrq1, resides in yeast. Both Hrq1 and Sgs1 are involved in genome integrity, functioning in processes such as DNA inter-strand crosslink repair, double-strand break repair, and telomere maintenance. However, it is unknown if these enzymes interact at a genetic, physical, or functional level as demonstrated for their human homologs. Thus, we performed synthetic genetic array (SGA) analyses of *hrq1Δ* and *sgs1Δ* mutants. As inactive alleles of helicases can demonstrate dominant phenotypes, we also performed SGA analyses on the *hrq1-K318A* and *sgs1-K706A* ATPase/helicase-null mutants, as well as all combinations of deletion and inactive double mutants. We crossed these eight query strains (*hrq1Δ, sgs1Δ, hrq1-K318A, sgs1-K706A, hrq1Δ sgs1Δ, hrq1Δ sgs1-K706A, hrq1-K318A sgs1Δ*, and *hrq1-K318A sgs1-K706A*) to the *S. cerevisiae* single gene deletion and temperature-sensitive allele collections to generate double and triple mutants and scored them for synthetic positive and negative genetic effects based on colony growth. These screens identified hundreds of synthetic interactions, supporting the known roles of Hrq1 and Sgs1 in DNA repair, as well as suggesting novel connections to rRNA processing, mitochondrial DNA maintenance, transcription, and lagging strand synthesis during DNA replication.

## INTRODUCTION

The human genome encodes five RecQ family helicases (RECQL1, BLM, WRN, RECQL4, and RECQL5), all of which are involved in the maintenance of genome integrity (Bochman 2014; Croteau *et al*. 2014). Two RecQ family helicases exist in *Saccharomyces cerevisiae*, Hrq1 and Sgs1, which are homologs of the disease-linked human RECQL4 (Barea *et al*. 2008; Bochman *et al*. 2014; Rogers *et al*. 2017) and BLM helicases (Watt *et al*. 1996; Lillard-Wetherell *et al*. 2005; Gravel *et al*. 2008), respectively. However, the discovery of Sgs1 (Gangloff *et al*. 1994) preceded that of Hrq1, and for many years, Sgs1 was considered the only RecQ family helicase encoded in the *S. cerevisiae* genome. However, a second DNA helicase with RecQ homology was independently identified several times (Shiratori *et al*. 1999; Lee *et al*. 2005), but Hrq1 was never formally named and recognized as a homolog of the RECQL4 helicase until 2008 (Barea *et al*. 2008), with *in vivo* and *in vitro* functional homology to RECQL4 being demonstrated subsequently (Bochman *et al*. 2014; Rogers and Bochman 2017; Rogers *et al*. 2017; Nickens *et al*. 2018; Rogers *et al*. 2020).

The known and hypothesized roles of Sgs1 in homologous recombination, DNA replication, meiosis, excision repair, and telomere maintenance were recently reviewed (Gupta and Schmidt 2020). Much less is known about Hrq1, though it is linked to DNA inter-strand crosslink (ICL) repair, telomere maintenance, and the unwinding of noncanonical DNA secondary structures (Bochman *et al*. 2014; Rogers and Bochman 2017; Rogers *et al*. 2017; Nickens *et al*. 2018; Rogers *et al*. 2020) like human RECQL4 (Jin *et al*. 2008; Ghosh *et al*. 2011; Ferrarelli *et al*. 2013; Keller *et al*. 2014). Contemporaneous work using a multi-omics approach also suggests that Hrq1 has roles in transcription, chromosome/chromatin dynamics, rRNA processing/ribosomal maturation, and in the mitochondria (Rogers et al.)^2^.

Despite these advances in yeast RecQ research, little is known about the genetic interactions that occur between *HRQ1* and *SGS1* or the physical interactions between Hrq1 and Sgs1. In humans, some of the RecQ helicases are partially functionally redundant (*e.g*., BLM and WRN), some display complementarity (*e.g*., WRN and RECQL5), and others exhibit functional synergism (reviewed in (Croteau *et al*. 2014)). The latter is exemplified by BLM and RECQL4, where BLM promotes the retention of RECQL4 at DNA double-strand breaks (DSBs), and RECQL4 stimulates BLM activity (Singh *et al*. 2012). Do such connections exist between their yeast homologs Hrq1 and Sgs1? Two reports demonstrate that various combinations of *hrq1* and *sgs1* alleles display differential responses to DNA damage compared to single mutants (Bochman *et al*. 2014; Rogers *et al*. 2020), suggesting that functional interactions among the RecQ helicases also exist in *S. cerevisiae*.

### Rationale for screen

The study of yeast RecQ homologs has greatly expanded our mechanistic understanding of how these enzymes function in various DNA repair pathways, but the interplay between Hrq1 and Sgs1 and their roles in other biological processes are not known. We sought to identify genes whose mutation affects the growth of *hrq1* and/or *sgs1* mutant cells. Because inactive alleles of DNA helicases often act as dominant negatives (Wu and Brosh 2010) and in some cases better represent disease-linked alleles, we utilized both deletion (*hrq1Δ* and *sgs1Δ*) and catalytically inactive mutants (*hrq1-K318A* and *sgs1-K706A*) of the helicases in all combinations (*hrq1Δ, sgs1Δ, hrq1-K318A, sgs1-K706A, hrq1Δ sgs1Δ, hrq1Δ sgs1-K706A, hrq1-K318A sgs1Δ*, and *hrq1-K318A sgs1-K706A*) in our screen. Many genes that encode proteins involved in genome integrity are also essential, so we performed synthetic genetic array (SGA) analysis by mating our query helicase mutant strains to both the *S. cerevisiae* single-gene deletion collection (Giaever and Nislow 2014) and the temperature-sensitive (TS) collection (Kofoed *et al*. 2015), the latter of which includes alleles of essential genes not found in the former, to generate a comprehensive set of double and triple mutant strains for SGA analysis.

## MATERIALS & METHODS

### Screen design

The strains used in this study are listed in Table 1. The *HRQ1* gene was deleted in Y8205 (Table 1) by transforming in a NatMX cassette that was PCR-amplified from the plasmid pAC372 (a gift from Amy Caudy) using oligonucleotides MB525 and MB526 (Table S1). The deletion was verified by PCR analysis using genomic DNA and oligonucleotides that anneal to regions up- and downstream of the *HRQ1* locus (MB527 and MB528). The confirmed *hrq1Δ* strain was named MBY639. The *hrq1-K318A* allele was introduced into the Y8205 background in a similar manner. First, an *hrq1-K318A(NatMX*) cassette was PCR-amplified from the genomic DNA of strain MBY346 (Bochman *et al*. 2014) using oligonucleotides MB527 and MB528 and transformed into Y8205. Then, genomic DNA was prepared from transformants and used for PCR analyses of the *HRQ1* locus with the same oligonucleotide set to confirm insertion of the NatMX marker. Finally, PCR products of the expected size for *hrq1-K318A(NatMX*) were sequenced using oligonucleotide MB932 to confirm the presence of the K318A mutation. The verified *hrq1-K318A* strain was named MBY644.

**Table 1.** Strains used in this study.

| Name | Genotype | Source |
| --- | --- | --- |
| Y8205 | *MATα can1Δ::STE2pr-Sp_his5 lyp1Δ::STE3pr-LEU2 his3Δ1 leu2Δ0 ura3Δ0* | (TONG *et al.* 2001) |
| MBY346 | *MATα ura3-52 lys2-801_amber ade2-101_ochre trp1Δ63 his3Δ200 leu2Δ1 hxt13::URA3 hrq1::hrq1-K318A-NatMX* | (BOCHMAN *et al.* 2014) |
| MBY639 | *MATα can1Δ::STE2pr-Sp_his5 lyp1Δ::STE3pr-LEU2 his3Δ1 leu2Δ0 ura3Δ0 hrq1::natMX* | This study |
| MBY640 | *MATα can1Δ::STE2pr-Sp_his5 lyp1Δ::STE3pr-LEU2 his3Δ1 leu2Δ0 ura3Δ0 sgs1::natMX* | This study |
| MBY642 | *MATα can1Δ::STE2pr-Sp_his5 lyp1Δ::STE3pr-LEU2 his3Δ1 leu2Δ0 ura3Δ0 sgs1::sgs1-K706A(natMX)* | This study |
| MBY643 | *MATα can1Δ::STE2pr-Sp_his5 lyp1Δ::STE3pr-LEU2 his3Δ1 leu2Δ0 ura3Δ0 hrq1::natMX sgs1::URA3* | This study |
| MBY644 | *MATα can1Δ::STE2pr-Sp_his5 lyp1Δ::STE3pr-LEU2 his3Δ1 leu2Δ0 ura3Δ0 hrq1::hrq1-K318A(natMX)* | This study |
| MBY645 | *MATα can1Δ::STE2pr-Sp_his5 lyp1Δ::STE3pr-LEU2 his3Δ1 leu2Δ0 ura3Δ0 hrq1::hrq1-K318A(natMX6) sgs1::URA3* | This study |
| MBY674 | *MATα can1Δ::STE2pr-Sp_his5 lyp1Δ::STE3pr-LEU2 his3Δ1 leu2Δ0 ura3Δ0 sgs1::sgs1-K706A(natMX6) hrq1::URA3* | This study |
| MBY676 | *MATα can1Δ::STE2pr-Sp_his5 lyp1Δ::STE3pr-LEU2 his3Δ1 leu2Δ0 ura3Δ0 hrq1::hrq1-K318A(URA3) sgs1::sgs1-K706A(natMX6)* | This study |

The *SGS1* gene was deleted from Y8205 (Table 1) in the same manner as the *HRQ1::natMX* deletion above by transforming in a NatMX cassette that was PCR-amplified using oligonucleotides MB1395 and MB768 (Table S1). The deletion was verified by PCR analysis of genomic DNA and oligonucleotides MB373 and MB374. The confirmed *sgs1Δ* strain was named MBY640. The *sgs1-K706A* allele was PCR amplified from plasmid pFB-MBP-Sgs1K706A-his (Cejka and Kowalczykowski 2010) (Table 2) using oligonucleotides MB765 and MB1396. The NatMX cassette was PCR-amplified from pAC372 using oligonucleotides MB1397 and MB768 and fused to the *sgs1-K706A* PCR product by Gibson assembly (Gibson *et al*. 2009). The resultant *sgs1-K706A*(*natMX*) cassette was reamplified with MB765 and MB768 and transformed into Y8205. Genomic DNA was then prepared from transformants and used for PCR analyses of the *SGS1* locus with oligonucleotides MB373 and MB374 to confirm insertion of the cassette. Finally, PCR products of the expected size were sequenced using oligonucleotide MB769 to confirm the presence of the K706A mutation. The verified *sgs1-K706A* strain was named MBY642.

**Table 2.** Results of the SGA analyses for all query strains crossed to the single-gene deletion collection.

| Query strain | No. negative genetic interactions | No. positive genetic interactions | Total |
| --- | --- | --- | --- |
| *hrq1Δ* | 76 | 41 | 117 |
| *hrq1-K318A* | 84 | 48 | 132 |
| *sgs1Δ* | 164 | 148 | 312 |
| *sgs1-K706A* | 189 | 172 | 361 |
| *hrq1Δ sgs1Δ* | 361 | 333 | 694 |
| *hrq1Δ sgs1-K706A* | 392 | 396 | 788 |
| *hrq1-K318A sgs1Δ* | 442 | 438 | 880 |
| *hrq1-K318A sgs1-K706A* | 400 | 396 | 796 |

The double mutant strains were constructed using similar techniques. Briefly, the *hrq1Δ sgs1Δ* and *hrq1-K318A sgs1Δ* strains were generated by deleting *SGS1* in strains MBY639 and MBY644, respectively, using a *URA3* cassette amplified from pUG72 (Gueldener *et al*. 2002) with oligonucleotides MB1395 and MB355 (Table S1). The strains verified by PCR of genomic DNA and sequencing were named MBY643 and MBY645, respectively. The *hrq1Δsgs1-K706A* and *hrq1-K318A sgs1-K706A* strains were constructed by amplifying *sgs1-K706A* as above, amplifying the *URA3* cassette with oligonucleotides MB1397 and MB355, and fusing the PCR products via Gibson assembly. The *sgs1-K706A(URA3*) cassette was then transformed into strains MBY639 and MBY644, and transformants were confirmed for proper integration by PCR and Sanger sequencing. The verified *hrq1Δ sgs1-K706A* and *hrq1-K318A sgs1-K706A* strains were named MBY674 and MBY676, respectively. Further details concerning strain construction are available upon request.

SGA analysis of the *hrq1Δ, sgs1Δ, hrq1-K318A, sgs1-K706A, hrq1Δ sgs1Δ, hrq1Δ sgs1-K706A, hrq1-K318A sgs1Δ*, and *hrq1-K318A sgs1-K706A* mutants was performed at the University of Toronto using previously described methods (Tong *et al*. 2001; Tong *et al*. 2004). All query strains and the control *HO::natMX* strain were crossed in quadruplicate to both the *S. cerevisiae* single-gene deletion collection (Giaever and Nislow 2014) and the TS alleles collection (Kofoed *et al*. 2015) to generate double or triple mutants for analysis.

### Phenotypes

Quantitative scoring of the genetic interactions was based on colony size. The SGA score measures the extent to which the size of a double or triple mutant colony differs from the colony size expected from combining the query and tester mutations together. The data includes both negative (putative synthetic sick/lethal) and positive interactions (potential epistatic or suppression interactions) (Tables S2-17). The magnitude of the SGA score is indicative of the strength of the interaction. Based on statistical analysis, it was determined that a default cutoff for a significant genetic interaction is *p* < 0.05 and SGA score > |0.08|.

### Verification of mutants

The top five negative and positive interactions for each query strain were confirmed by remaking and reanalyzing the double and triple mutants by hand, followed by spot dilution assays to compare the growth of the double or triple mutants to their parental strains and wild-type.

### Statistical analysis

Data were analyzed and graphed using GraphPad Prism 6 software. The reported values are averages of ≥ 3 independent experiments, and the error bars are the standard deviation. *P*-values were calculated as described in the figure legends, and we defined statistical significance as *p* < 0.01.

### Data availability

Strains, plasmids, and other experimental reagents are available upon request. File S1 contains Table S1 and a description of the other supplementary tables included in Files S2 and S3. File S2 contains Tables S2-S9, and File S3 contains Tables S10-S17.

## RESULTS AND DISCUSSION

### Overall results of the screen

Hundreds of synthetic interactions were detected for all query strains screened through both the single-gene deletion (Table 2) and TS mutant (Table 3) collections (Tables S2-17). For the single-gene deletion collection screen, the numbers of negative and positive genetic interactions were generally the same for all query strains, except *hrq1Δ* and *hrq1-K318A*, which yielded approximately twice as many negative as positive interactions (Table 2). These mutants also had the fewest number of synthetic interactions by a factor of > 2.3 compared to *sgs1Δ* and *sgs1-K706A*. This is consistent with the generally more modest phenotypes of *hrq1Δ* and *hrq1-K318A* strains compared to *sgs1Δ* and *sgs1-K706A* for DNA damage sensitivity (Bochman *et al*. 2014). The double mutant query strains yielded a greater than additive number of synthetic genetic interactions than the single mutant parental query strains, indicating that mutating both RecQ helicases had a synergistic effect. This synergism was strongest for the *hrq1-K318A sgs1Δ* mutant, which generated 880 synthetic interactions, a nearly twofold increase over the additive effect of the 132 *hrq1-K318A* and 312 *sgs1Δ* interactions individually (compared to ~1.5- to 1.6-fold increases for the other combinations).

**Table 3.** Results of the SGA analyses for all query strains crossed to the temperature-sensitive allele collection.

| Query strain | No. negative genetic interactions | No. positive genetic interactions | Total |
| --- | --- | --- | --- |
| *hrq1Δ* | 65 | 54 | 119 |
| *hrq1-K318A* | 82 | 61 | 143 |
| *sgs1Δ* | 155 | 197 | 352 |
| *sgs1-K706A* | 138 | 172 | 310 |
| *hrq1Δ sgs1Δ* | 156 | 246 | 402 |
| *hrq1Δ sgs1-K706A* | 238 | 260 | 498 |
| *hrq1-K318A sgs1Δ* | 200 | 268 | 468 |
| *hrq1-K318A sgs1-K706A* | 223 | 232 | 455 |

For the TS allele collection screen, the numbers of negative and positive genetic interactions were again generally similar for all query strains (Table 3). As above, the *hrq1Δ* and *hrq1-K318A* mutants had the fewest number of synthetic interactions by a factor of > 2.1 compared to *sgs1Δ* and *sgs1-K706A*. In this case, however, the double mutant query strains yielded approximately an additive number of synthetic genetic interactions compared to the single mutant parental query strains and thus did not display the synergism described for the single-gene deletion SGA analysis. It should also be noted that the numbers of synthetic genetic interactions listed in Table 3 are inflated because several different TS alleles of the same ORF are included in the collection for many individual genes (Kofoed *et al*. 2015).

Figure 1 shows the frequency distribution of all of the SGA scores as violin plots and separate box plots of the negative and positive synthetic genetic interactions, with outliers denoted as single points, for the single-gene deletion collection (Fig. 1A-C) and the TS collection (Fig. 1D-F). The outliers represent the mutants with the strongest synthetic phenotypes. As shown in Figures 1A and 1D, most synthetic phenotypes were mild decreases or increases in the growth of the double and triple mutant colonies. There were no significant differences in the distribution of the SGA scores among any of the mutant sets generated by crossing the query strains to the single-gene deletion collection. However, several significant differences were found in the distributions of positive SGA scores for the mutant sets yielded from the crosses to the TS collection. These includes mild differences between *hrq1-K318A vs. hrq1Δ sgs1Δ (p* = 0.0123) and *sgs1Δ vs. hrq1-K318A sgs1Δ (p* = 0.0303), intermediate differences for *sgs1Δ vs. hrq1Δ sgs1Δ (p* = 0.0016) and *sgs1-K706A vs. hrq1-K318A sgs1-K706A (p* = 0.0070), and strong differences between *sgs1-K706A* and *hrq1Δ sgs1Δ, hrq1Δ sgs1-K706A*, and *hrq1-K318A sgs1Δ* (all *p* < 0.0001). It is currently unclear why the strength of the positive synthetic genetic interactions significantly varied among these mutants, especially compared to the *sgs1-K706A* query strain, but we are actively following up on phenotypic difference among all of the *hrq1* and *sgs1* alleles. Regardless, as mutants giving the strongest growth effects, the outliers in Figures 1B, C, E, and F are summarized in Tables 4 and 5. For simplicity, only the negative genetic interactions are discussed in further detail below.

**Table 4.**
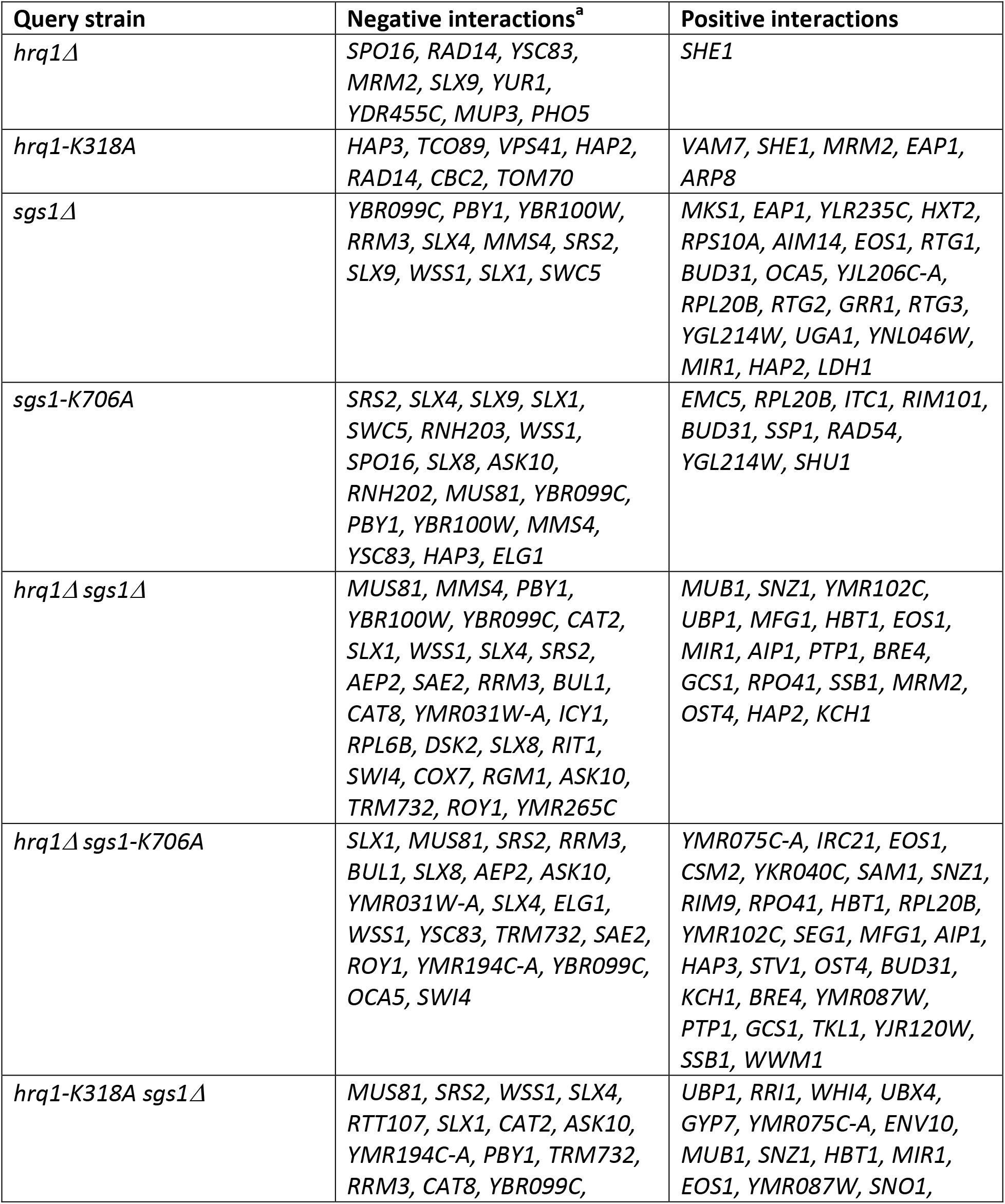
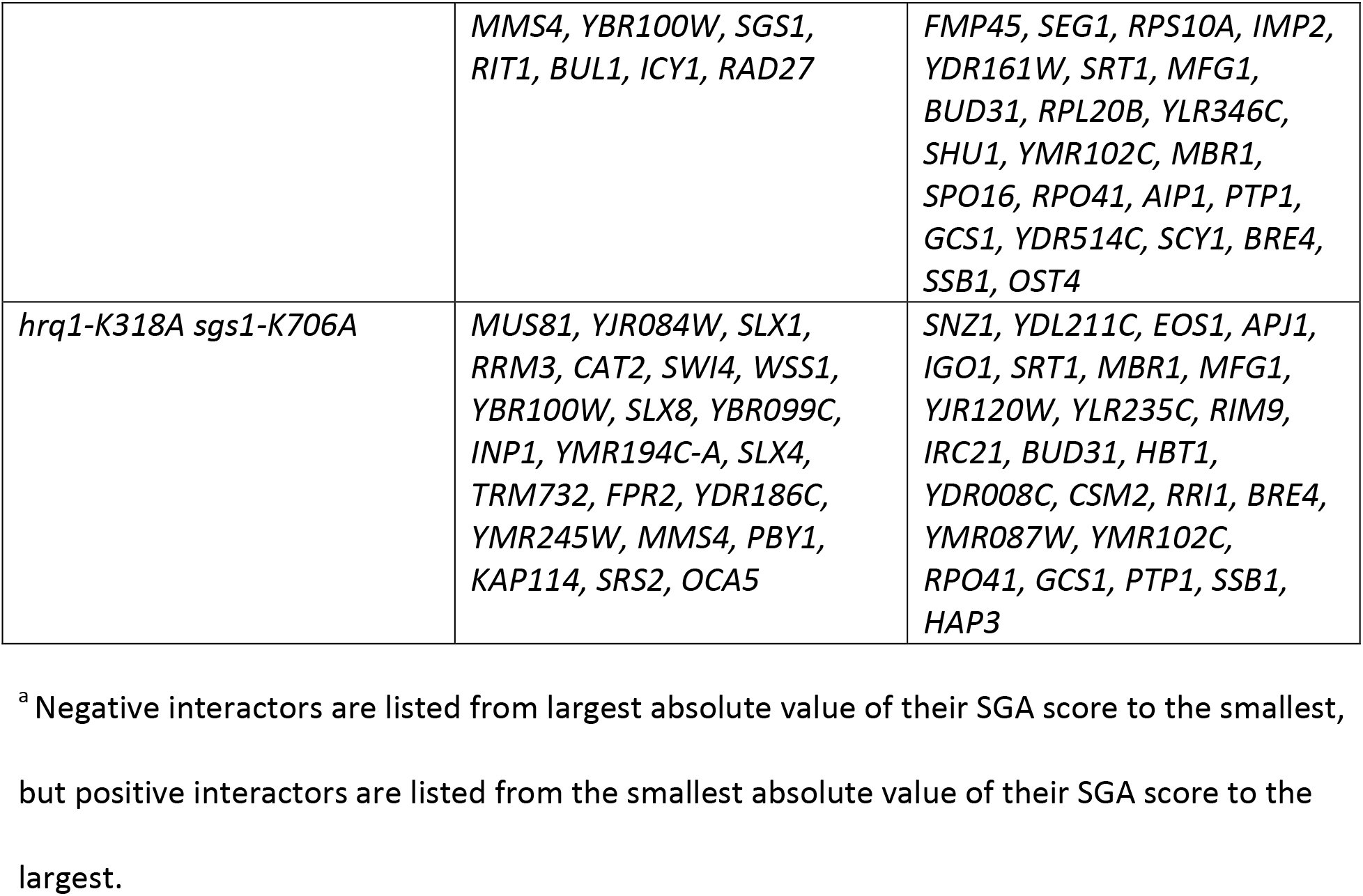
Genes whose deletion cause the strongest growth phenotypes when combined with the *hrq1* and *sgs1* mutants.

**Table 5.** Temperature-sensitive alleles that cause the strongest growth phenotypes when combined with the *hrq1* and *sgs1* mutants.

| Query strain | Negative interactions<sup>a</sup> | Positive interactions |
| --- | --- | --- |
| *hrq1*Δ | *yef3-f650s* | *mps3-1, slx9, act1-105, arp3-g302y, arp3-31, crm1-1* |
| *hrq1-K318A* | *yef3-f650s* | *cse2, dbp5-2, crm1-1, mps1-1* |
| *sgs1*Δ | *yef3-f650s, efb1-4, swc4-4* | *stt3-1, arp2-14, ydl003w-ph, sfi1-7, nse5-ts1, rad54, nsl1-6, act1-105, mps3-1, smc2-8, hrp1-7, crm1-1* |
| *sgs1-K706A* | *ypr086w-ph, yef3-f650s, swc4-4* | *brn1-9, ydr331w-ph, mps3-1, arc40-ph, cdc20-3, dpb11-1, arp3-g302y, crm1-1* |
| *hrq1*Δ *sgs1*Δ | *dna2-1, nse4-ts2, mob2-38, smt3-331, rot1-ph, mob2-22* | *kch1, pol1-17, nuf2-ph, pse1-41, ndc1-4, tor2-29, mps1-1, nsl1-6, prp6-ts, ypr086w-ph, mps3-1, rad5* |
| *hrq1*Δ *sgs1-K706A* | *nse4-ts2, dbf4-1, rot1-ph, cdc2-1, mob2-28, cep3-1, nse4-ts1, mob2-22, smc6-9, mob2-38, mob2-14, pri2-1, nse1-16, mob2-8* | *arf1, ala1-1, gna1-ts, gle1-4, sfi1-7, nut1, tim22-19, prp6-ts, cdc23-1, cdc20-1, cdc13-1, rad54, nse5-ts1, yol102c-ph* |
| *hrq1-K318A sgs1*Δ | *dbf4-1, dna2-1, sgs1, nse4-ts2* | *yhr122w-ph, nut1, sfi1-7, pse1-41, yjl174w-ph, cab1-ph, srm1-ts, kch1, tim22-19, sth1-3, yjl011c-ph, prp6-ts, bem1, yol102c-ph, act1-105* |
| *hrq1-K318A sgs1-K706A* | *nse4-ts2, rot1-ph, cdc2-1, mob2-38* | *lip1-ph, ala1-1, gle1-4, srv2-2, sfh1-1, nut1, pse1-41, sth1-3, cdc28-td, gpi17-ph, tim22-19, arp3-31, yol102c-ph, prp43-ts2* |

<sup>a</sup> Negative interactors are listed from largest absolute value of their SGA score to the smallest, but positive interactors are listed from the smallest absolute value of their SGA score to the largest.

**Figure 1.**
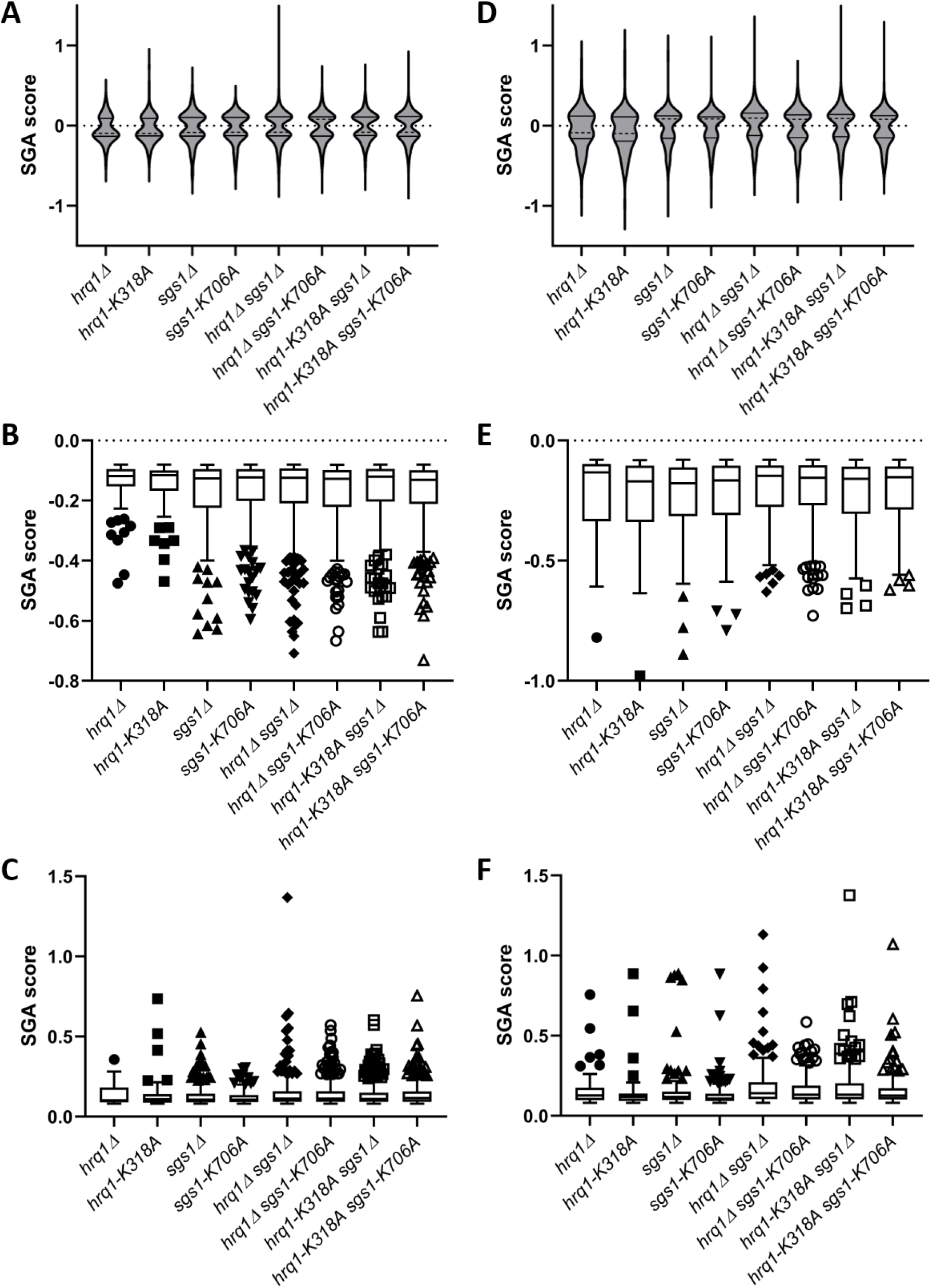
Analysis of the distribution of the magnitudes of the synthetic genetic interactions. Violin plots of the synthetic genetic interactions with the single-gene deletion collection (A) and TS collection (D). The median values are denoted with dashed lines, and the quartiles are shown as solid lines. The SGA data are also shown in separate box and whisker plots drawn using the Tukey method for the negative (B) and positive (C) interactions with the deletion collection, as well as for the negative (E) and positive (F) interactions with the TS collection. The individually plotted points outside of the inner fences represent outliers (*i.e*., interactions with mutants yielding the strongest SGA scores) and correspond to alleles whose SGA score is less than the value of the 25^th^ quartile minus 1.5 times the inter-quartile distance (IQR) for negative interactions and alleles whose SGA score is greater than the value of the 75^th^ quartile plus 1.5IQR for positive interactions. The significant differences between SGA data sets discussed in the main text were calculated using the Kruskal-Wallis test and Dunn’s multiple comparisons test.

### *hrq1Δ* interactions

The deletion of *HRQ1* displayed strong negative interactions with mutations in 10 genes (Tables

4 and 5), many of which correspond to the recently described Hrq1 interactome (Rogers *et al*.)^1^. For instance, RECQL4 is the only human RecQ found in both the nucleus and mitochondria (Croteau *et al*. 2014), and Hrq1 likewise localizes to both organelles (Koh *et al*. 2015) and physically interacts with mitochondrial proteins (Rogers *et al*.)^1^. Here, we found strong negative synthetic genetic interactions between *hrq1Δ* and mutation of *MRM2*, a mitochondrial 2’ O-ribose methyltransferase whose deletion results in mitochondrial DNA (mtDNA) loss (Pintard *et al*. 2002), and *YSC83*, a mitochondrial protein of unknown function (Sickmann *et al*. 2003). It is still unclear what the role of Hrq1 is in the mitochondria, but it is tempting to speculate that it is involved in mtDNA maintenance in a similar fashion to its maintenance of the nuclear genome.

This role in genome integrity is highlighted by the negative interactions of *hrq1Δ* with mutation of *SPO16*, which is involved in the meiotic cell cycle (Shinohara *et al*. 2008), and *RAD14*, a nucleotide excision repair protein (Guzder *et al*. 2006) and regulator of transcription (Chaurasia *et al*. 2013). Deletion of *HRQ1* also negatively interacted with mutation of *SLX9*, an rRNA processing factor (Bax *et al*. 2006) that additionally binds G-quadruplex (G4) DNA structures (Gotz *et al*. 2019). This is provocative in light of the connection of Hrq1 to rRNA processing and ribosome biogenesis (Rogers *et al*.)^1^, as well as the fact that G4 structures are preferred substrates for Hrq1 *in vitro* (Rogers *et al*. 2017). Finally, mutations in *YEF3, YUR1, MUP3*, and *PHO5* (encoding a translation elongation factor, protein glycosylase, methionine permease, and acid phosphatase, respectively), as well as the dubious open reading frame (ORF) *YDR455C* (Fisk *et al*. 2006), also negatively interacted with *hrq1Δ*.

### *hrq1-K318A* interactions

Mutations in only two genes, *RAD14* and *YEF3*, are shared between the lists of strong negative interactors with *hrq1Δ* and *hrq1-K318A*. This is not unexpected based on the ability of Hrq1-K318A to phenocopy wild-type in some pathways (Bochman *et al*. 2014). However, mutations in genes encoding proteins involved in processes shared between both sets are evident. This includes *HAP2* and *HAP3*, which are activators of transcription (Xing *et al*. 1993), *TCO89*, a member of the TOR complex and global regulator of histone H3 K56 acetylation (Chen *et al*. 2012), and *RAD14* as described above. Similarly, *TOM70* encodes a subunit of the mitochondrial protein importer (Brix *et al*. 2000), which is likely important for localizing Hrq1 to the mitochondria where it may be involved in mtDNA maintenance. Genome integrity is also highlighted by *CBC2*, which encodes an RNA binding and processing factor involved in telomere maintenance (Lee-Soety *et al*. 2012). Hrq1 is known to regulate telomerase activity at both DSBs and telomeres (Bochman *et al*. 2014; Nickens *et al*. 2018; Nickens *et al*. 2019). Mutation of the gene encoding the Vps41 vacuolar membrane protein (Nakamura *et al*. 1997) also negatively interacted with *hrq1-K318A*.

### *sgs1Δ* interactions

Over 500 genetic interactions with *sgs1* alleles have been reported (see: https://www.yeastgenome.org/locus/S000004802/interaction), including most of the hits from our screen, such as the genome integrity genes *MMS4, RRM3, SLX1, SLX4, SRS2*, and *WSS1* (Fisk *et al*. 2006), as well as *SLX9* (see above) and *EFB1*, which encodes a translation elongation factor (Hiraga *et al*. 1993). These hits serve as internal positive controls. It should also be noted that: 1) *YBR099C* is a dubious ORF that completely overlaps *MMS4* (Fisk *et al*. 2006), 2) *YBR100W* was an originally misannotated ORF and more recently merged with an adjacent ORF such that the coding region is now named *MMS4* (Xiao *et al*. 1998), and 3) the *pby1Δ* strain in the single-gene deletion collection is actually a deletion of *MMS4* (Olmezer et al. 2015). Thus, multiple different *mms4* alleles were hits in the screen, again acting as positive controls for our approach.

In addition to known effects, we also discovered three new negative interactions with *sgs1Δ*. These include the deletions of *SWC4* and *SWC5*, which encode subunits of the SWR1 complex that replaces histone H2A with H2A.Z (Mizuguchi *et al*. 2004), preventing the spread of silent heterochromatin (Meneghini *et al*. 2003). This interaction could be connected to the role of Sgs1 in telomere maintenance (Huang *et al*. 2001; Johnson *et al*. 2001; Azam *et al*. 2006) because telomeric DNA is also silenced via the telomere position effect (Mondoux and Zakian 2005). As with *hrq1Δ* and *hrq1-K318A*, the *yef3-f650s* TS allele was also a negative genetic interactor with *sgs1Δ* (Table 5).

### *sgs1-K706A* interactions

Unlike *sgs1Δ*, much less is known about the genetic interactome of the catalytically inactive *sgs1-K706A* allele. We found that the strong negative interactors were mutations in genes that completely overlap with the *sgs1Δ* set (*SRS2, SLX4*, SLX9, *SLX1, SWC5, WSS1, MMS4, ELG1, YEF3*, and *SWC4*). However, the *sgs1-K706A* interactors also included mutations in genes that were not ranked as causing the strongest negative effects with *sgs1Δ*. Nevertheless, alleles of some of these genes (*RNH203, SLX8, RNH202*, and *MUS81*) are previously reported negative interactors with *sgs1Δ* (see: https://www.yeastgenome.org/locus/S000004802/interaction).

Mutations in the remaining genes have not previously been reported to negatively interact with *sgs1Δ*, but three of them (*SPO16, YSC83*, and *HAP3*) overlap with the *hrq1* interactors described above, perhaps suggesting some overlap in function between Hrq1 and Sgs1 in the pathways related to these genes. That leaves only two genes, *SUA7* and *ASK10*, as unique interactors here. The *SUA7* gene product is the yeast transcription factor TFIIB that is needed for RNA polymerase II transcriptional start site selection (Pinto *et al*. 1992). This may indicate that like the human RECQL5 helicase (Aygun *et al*. 2008; Izumikawa *et al*. 2008; Saponaro *et al*. 2014), Sgs1 is involved in transcription, a hypothesis also put forth for Hrq1 (Rogers *et al*.)^1^. In support of this, the remaining interactor *ASK10* encodes a glycerol channel regulator (Beese *et al*. 2009) that also associates with RNA polymerase II (Page *et al*. 1996).

### Negative genetic interactions with the *hrq1 sgs1* double mutants

The sets of synthetic negative genetic interactions for the *hrq1 sgs1* double mutants shown in Tables 4 and 5 generally contain the strong interactors from the single-mutant parental strains, but they also include many new interactions, evident of the synergistic effect of mutating both RecQ family helicases in *S. cerevisiae*. These genes (*CAT2, AEP2, SAE2, BUL1, CAT8, YMR031W-A, ICY1, RPL6B, DSK2, RIT1, SWI4, COX7, RGM1, TRM732, ROY1, YMR265C, ELG1, YMR194C-A, OCA5, RTT107, RAD27, YJR084W, INP1, FPR2, YDR186C, YMR245W, KAP114, DNA2, NSE4, MOB2, SMT3, ROT1, DBF4, CDC2, CEP3, SMC6, PRI2*, and *NSE1*) are enriched for gene ontology terms related to genome integrity, including DNA repair (*DNA2, ELG1, NSE1, NSE4, POL3, PRI2, RAD27, RTT107, SAE2*, and *SMC6*), DNA replication (*DBF4, DNA2, ELG1, CDC2, PRI2*, and *RAD27*), and transcription by RNA polymerase II (*CAT8, CEP3, RGM1, SWI4*, and *YJR084W*) among others.

The links to DNA replication are notable because the negative genetic interactions preferentially occur with genes encoding lagging strand synthesis machinery: Dna2 and Rad27 are both nucleases involved in Okazaki fragment processing (Kao *et al*. 2004), Cdc2 is the catalytic subunit of DNA polymerase δ (Johnson *et al*. 2015), and Pri2 is the large subunit of DNA primase (Foiani *et al*. 1989). It is also known that both Hrq1 (Bochman *et al*. 2014; Nickens *et al*. 2018) and Sgs1 (Wagner *et al*. 2006) interact with the Pif1 helicase, an enzyme involved in the two-nuclease Okazaki fragment processing pathway (Rossi *et al*. 2008; Pike *et al*. 2009). Therefore, combinatorial mutations of both yeast RecQ helicases are strongly deleterious when lagging strand synthesis is also disrupted by mutation. It is tempting to speculate that hindered Okazaki fragment maturation may yield DNA structures or lesions that require the repair activities of Hrq1 and Sgs1 for processing.

Also intriguing are the genes of unknown function (*YMR265C, ICY1*, and *YMR245W*) and those categorized as dubious ORFs (*YMR194C-A* and *YMR031W-A*) (Fisk *et al*. 2006). For instance, even though it is a dubious ORF, deletion of *YMR031W-A* yields cells with short telomeres (Askree *et al*. 2004), and Hrq1 (Bochman *et al*. 2014; Rogers *et al*. 2017; Nickens *et al*. 2018) and Sgs1 (Watt *et al*. 1996; Huang *et al*. 2001; Johnson *et al*. 2001; Azam *et al*. 2006) are both involved in telomere maintenance. Further research should be devoted to uncovering the links between the *YMR265C, ICY1, YMR245W, YMR194C-A*, and *YMR031W-A* gene products and RecQ biology in *S. cerevisiae*.

### Conclusions and perspectives

Here, we have reported a comprehensive set of synthetic genetic interactions between most of the genes in the *S. cerevisiae* genome and deletion and catalytically inactive alleles of the Hrq1 and Sgs1 RecQ family helicases. This data set improves upon the existing sets of known *hrq1Δ* and *sgs1Δ* interactions and expands the genetic interactome landscape of *hrq1* and *sgs1* mutants by including interactions with the inactive *hrq1-K318A* and *sgs1-K706A* alleles, as well as all combinations of the null and inactive double mutants. As with the five human RecQ helicases (Croteau *et al*. 2014), it is clear that *HRQ1* and *SGS1* genetically interact in yeast, and perhaps they may also physically interact.

These SGA analyses have also generated testable hypotheses to drive on-going and future research. The genetic interactomes of *hrq1* and *sgs1* suggest links to transcription, much like the functional interaction between human RECQL5 and RNA polymerase II (Aygun *et al*. 2008; Izumikawa *et al*. 2008; Saponaro *et al*. 2014). Indeed, we have already shown that *hrq1* cells are sensitive to the general transcription inhibitor caffeine and that *hrq1* mutations alter the *S. cerevisiae* transcriptome (Rogers *et al*.)^1^. Similarly, it will be exciting to discover why double *hrq1 sgs1* mutations are particularly deleterious to defects in lagging strand synthesis during DNA replication.

Obviously, our focus on the strongest negative synthetic genetic interactions in the SGA data set reported here is far from all encompassing. There are certainly important conclusions to be drawn from more subtle negative effects, considering the positive genetic interactions, and comparing the genetic interactomes between the various *hrq1* and *sgs1* mutants analyzed. It is our hope that these data will spur additional research in the field, both with the yeast RecQs and their human homologs RECQL4 and BLM, as well as with proteomic investigations to incorporate physical interactomes, to fully establish the roles of these enzymes in genome integrity.

## ACKNOWLEDGEMENTS

We thank Amy Caudy and Stephen Kowalczykowski for sharing plasmids, the University of Toronto for performing the SGA analyses, Michael Costanzo and members of the Boone lab for help with data collection and interpretation, and members of the Bochman lab for critically reading this manuscript. This research was supported by the College of Arts and Sciences, Indiana University (to MLB), the Indiana University Collaborative Research Grant fund of the Office of the Vice President for Research (to MLB), the American Cancer Society (RSG-16-180-01-DMC to MLB), and the National Institutes of Health (1R35GM133437 to MLB).

## FOOTNOTES

1 Rogers *et al*., The genetic and physical interactomes of the *Saccharomyces cerevisiae* Hrq1 helicase, <u>submitted as a companion paper to *G3*</u>.

2 Rogers et al., The genetic and physical interactomes of the *Saccharomyces cerevisiae* Hrq1 helicase, <u>submitted as a companion paper to *G3*</u>.

